# CD68^+^ Follicular Macrophages Harbor HIV Reservoirs in Human Lymph Node Tissues During Suppressive ART

**DOI:** 10.1101/2025.02.03.636184

**Authors:** Merantha Moodley, Caroline Chasara, Trevor Khaba, Bongiwe Mahlobo, Nicole Reddy, Sifundo Nxele, Sibongiseni Msipa, Kavidha Reddy, Thandekile Ngubane, Ismail Jajbhay, Johan Pansegrouw, Thumbi Ndung’u, Zaza M. Ndhlovu

## Abstract

Uncertainty persists regarding the contribution of tissue macrophages to HIV reservoirs, largely due to insufficient characterization of these reservoirs within their native tissue microenvironments. This study aimed to characterize and quantify macrophage reservoirs in human lymph node (LN) tissues in terms of their phenotype, location, and their potential for sustained productive infection during suppressive antiretroviral therapy.

We examined the topology, nature, and size of macrophage reservoirs in lymph nodes (LNs) from 45 PLWH subtype C on suppressive ART and 14 matched controls using *in situ* imaging and multiplexed immunofluorescence microscopy. Germinal center CD68^+^ macrophages harbored HIV *gag-pol* DNA, HIV *gag-pol* RNA and Gagp24 protein. Digital droplet PCR confirmed the presence of proviral reservoirs in myeloid cells within LNs. High-resolution imaging techniques revealed that infected macrophages within GCs displayed distinct morphological characteristics, featuring larger and irregular shapes. In contrast, phagocytic macrophages exhibited intracellular staining for CD4^+^ T cells, had regular shapes, and were predominantly found outside the GCs.

Our findings provide detailed quantitative, spatial, and phenotypic characterization of macrophage reservoirs in LNs, offering a clear estimation of the extent to which macrophages contribute to persistent HIV reservoirs in these tissues. These findings establish a basis for developing targeted strategies aimed at the elimination of these reservoirs in LN tissues.

**Author summary:** HIV hides in reservoirs within immune cells across the body in the blood and various tissues, making it a complex challenge to cure. Therefore, it is essential to identify and understand all sources of HIV reservoirs aid the development of an HIV cure. Macrophages are increasingly recognized as key contributors of viral reservoir persistence. However, the role of macrophages as latent HIV reservoirs remains unclear due to limited studies in human tissues. In this study we investigated macrophage reservoirs in human lymph nodes to answer key questions: where do they hide, how can we identify them, what is their contribution to the lymph node reservoir burden, and are they truly able to support productive viral replication? We found that macrophages residing within lymph node germinal centers, identified by CD68 expression, contained HIV DNA, RNA, and Gagp24 protein. Moreover, using high resolution microscopy, we were able to distinguish between productively infected macrophages from those that engulfed T cells. By unveiling these unique features of macrophage reservoirs, our research paves the way for the design of targeted therapy aimed at eliminating these reservoirs, towards an HIV cure.

## Introduction

Comprehensive characterization of HIV reservoirs is essential for the success of curative strategies (1). Lymph node (LN) tissues serve as key sanctuary sites for HIV reservoir persistence during suppressive antiretroviral therapy (ART). Poor ART penetration and suboptimal immune responses pose major challenges to reservoir eradication in LNs (2). Although very early ART initiation has been shown to accelerate HIV reservoir decay low level HIV replication persists in secondary lymphoid organs (3, 4). We previously demonstrated that T follicular helper (Tfh) cells within germinal centers (GCs) are the primary reservoirs in LN tissues of individuals on ART who began therapy during hyperacute HIV infection (5). We also identified that insufficient trafficking of cytotoxic CD8^+^ T cells into GCs, caused by a lack of CXCR5 expression, as a major obstacle to immune-mediated HIV reservoir eradication in LNs (6).

In our previous research we observed that CD4^+^ T cells and follicular dendritic cells alone could not account for all reservoir-harboring cells within B cell follicles (BCFs), we hypothesized that myeloid cells, specifically lymphoid tissue-resident macrophages, could play a role in maintaining persistent HIV reservoirs in LN tissues (1, 5). This hypothesis is based on emerging strong evidence showing the ability of tissue macrophages to harbor productively HIV infected viral reservoirs despite suppressive ART in the urethra (7), and because macrophages also reside in niches where HIV reservoirs persist such as the GCs of LN tissues (8–11).

Macrophages are highly diverse, with distinct phenotypes differing in their contributions to HIV reservoirs (11, 12). This variability is influenced by their physiological functions and spatial localization within the microenvironment. Macrophage permissiveness to HIV replication, combined with the prolonged survival of certain subsets suggest they may serve as long-lived HIV reservoirs (13). The classification of macrophages into pro-inflammatory or anti-inflammatory states is complex, as a variety of markers can be used to distinguish these phenotypes such as: CD68^+^/CD80^+^/CD86^+^ (pro-inflammatory) or CD206^+^/CD163^+^/CD200^+^ (anti-inflammatory) (14–18). Common classification strategies to delineate macrophage subtypes involve the use of CD68 as a pan macrophage marker and CD206 to characterize anti-inflammatory macrophages (7, 14, 15). The identification of long-lived, embryonically-derived macrophages, combined with their inherent resistance to cytotoxic T lymphocyte (CTL)-mediated clearance, further emphasizes their significance as a stable HIV reservoir (7, 12, 13, 19–26).

While cumulative research has established the existence of replication-competent, long-lived HIV reservoirs in macrophages (11, 27, 28), the precise mechanisms and extent of their involvement in sustaining viral persistence remain contentious. Further investigation into macrophage reservoirs within secondary lymphoid tissues, known to harbor a higher HIV reservoir burden, is crucial for elucidating their specific role in HIV reservoir dynamics. This study utilized excisional LN tissue samples collected from 59 participants, 45 people living with HIV (PLWH) and 14 demographically matched people living without HIV (PLWoH). 60.00% of PLWH were virally suppressed, whereas 37.77% exhibited low but detectable viremia at the time of sample collection. The samples were analyzed using *in situ* hybridization and quantitative high-resolution multicolor immunofluorescence (IF) techniques. HIV infection was associated with a significant increase in the macrophage population within LN tissues. CD68^+^ macrophages residing in BCFs were found to harbor HIV proviral DNA, *gag-pol* RNA, and Gagp24 protein. Analysis of myeloid cells by digital droplet PCR (ddPCR) confirmed presence of proviral HIV DNA in 70% of the samples analyzed from PLWH. Importantly, high-resolution microscopy revealed distinct morphological differences between productively infected macrophages and those that may phagocytose CD4^+^ T cells. These findings underscore the critical role of macrophages as a reservoir for HIV in LNs during suppressive ART, highlighting the need to include this subset in future HIV eradication research aimed at achieving a cure.

## Results

### Participant demographics

We recruited 59 participants from the Females Rising through Education Support and Health (FRESH) and HIV Pathogenesis Programme (HPP) Acute study cohorts encompassing PLWoH (n=14), individuals treated during Fiebig stage I-V, here referred to as acute-treated PLWH (n=10) and individuals treated after Fiebig stage V, here referred to as chronic-treated PLWH (n=31). 4 PLWH had an unknown treatment status at the time of sample collection. The demographic and clinical characteristics of the participants including sex, age, LN site, plasma viral load (pVL), CD4^+^ T cell count, ART timing and ART duration prior to LN excision are presented in **(Table 1, see also S1 Table)**. The pVL was detectable in 2/10 (20%) acute-treated and 15/31 (48.38%) chronic-treated PLWH (1700 vs. 2576 mean cps/mL) **(S1 Table)**. The median days to ART initiation following HIV detection was 1 day in the acute-treated PLWH group; and only one chronic-treated individual had a documented HIV detection date which enabled a single value calculation of 310 days until ART initiation post-initial HIV detection **(Table 1)**.

**Table 1.** Participant demographics and clinical data.

|  | PLWoH<br>n=14 | Acute<br>Treated<br>PLWH<br>n=10 | Chronic<br>Treated<br>PLWH<br>n=31 | Unknown<br>treatment<br>status PLWH<br>n=4 | p |
| --- | --- | --- | --- | --- | --- |
| Sex female/male (% female)<sup>a</sup> | 14/0<br>(100%) | 10/0<br>(100%) | 27/4<br>(87%) | 3/1<br>(75%) | 0.1932 |
| Age, median (IQR)<sup>b</sup> | 21.50<br>(3.250) | 23.00<br>(4.250) | 23.00<br>(7.250) | 44.00<br>(23.00) | 0.0038 |
| Lymph node site inguinal/axillary or unknown site (%inguinal)<sup>a</sup> | 11/3<br>(78.5%) | 10/0<br>(100%) | 23/8<br>(74.2%) | 0/4<br>(0%) | 0.0026 |
| CD4 count, mean cells/L (SD)<sup>c</sup> | 857.3<br>(196.1) | 816.7<br>(230.3) | 621.5<br>(262.5) | 666.3<br>(262.1) | 0.0282 |
| Plasma viral load, median RNA copies/mL (IQR)<sup>b</sup> | N/A | <20<br>(397.5) | <20<br>(2290) | <20<br>(0) | 0.1191 |
| ART initiation, median days after HIV detection (IQR) | N/A | 1.000<br>(0.250) | 310<br>(*IQR) | - | N/A |
| ART duration, median days from ART initiation to lymph node excision (IQR)<sup>d</sup> | N/A | 421.0<br>(773.7) | 151.0<br>(868.0) | - | 0.9420 |

<sup>a</sup>: Fisher's exact test; <sup>b</sup>: Kruskal-Wallis test; <sup>c</sup>: One-way ANOVA; <sup>d</sup>: Mann-Whitney U Test; - Data not reported Other: \*IQR: Cannot calculate IQR – only 1 datapoint. IQR: Inter-quartile range and SD: standard deviation. See also Table S1.

### Impact of HIV infection on the phenotypes, location and frequencies of macrophages in human LN tissues

LN GCs are known to be a key anatomical site of HIV reservoir persistence (5, 29). While macrophages are prevalent in this microanatomical site and may play a role in facilitating HIV infection, the precise phenotypic characteristics and spatial distribution of macrophages susceptible to HIV infection have not been comprehensively characterized in human LNs (30). Given that LN GCs are HIV reservoir hotspots, we hypothesized that specific macrophage subsets residing within GCs significantly contribute to the establishment of these reservoirs. To test this, we initially performed multicolor IF microscopy to phenotype macrophages in both follicular and extrafollicular (EF) regions. We characterized pro-inflammatory macrophages as CD68^+^CD206^-^ and anti-inflammatory macrophages as CD68^+^CD206^+^ (7, 14, 15). The transcription factor BCL-6 was used to identify active GCs. As shown by representative images, we observed a distinct spatial distribution of macrophage subsets whereby the follicular regions mostly harbored CD68^+^CD206^-^ macrophages **(Fig 1A and S1 Fig)**, whereas CD68^+^CD206^+^ macrophages were mostly localized in EF regions, close to lymph and blood vessels **(Fig 1B and S1 Fig)**. CD68^+^ cells were confirmed as true macrophages by co-staining with an additional macrophage marker Iba-1 and demonstrating negative expression for CD3 (7) **(S1 Fig)**. Quantitative image analysis using TissueQuest **(Fig S2)** confirmed that the density (cells/mm²) of CD68⁺ macrophages was significantly higher within the EF regions than in GCs (p=0.0411) **(Fig 1C).**

**Fig 1.**
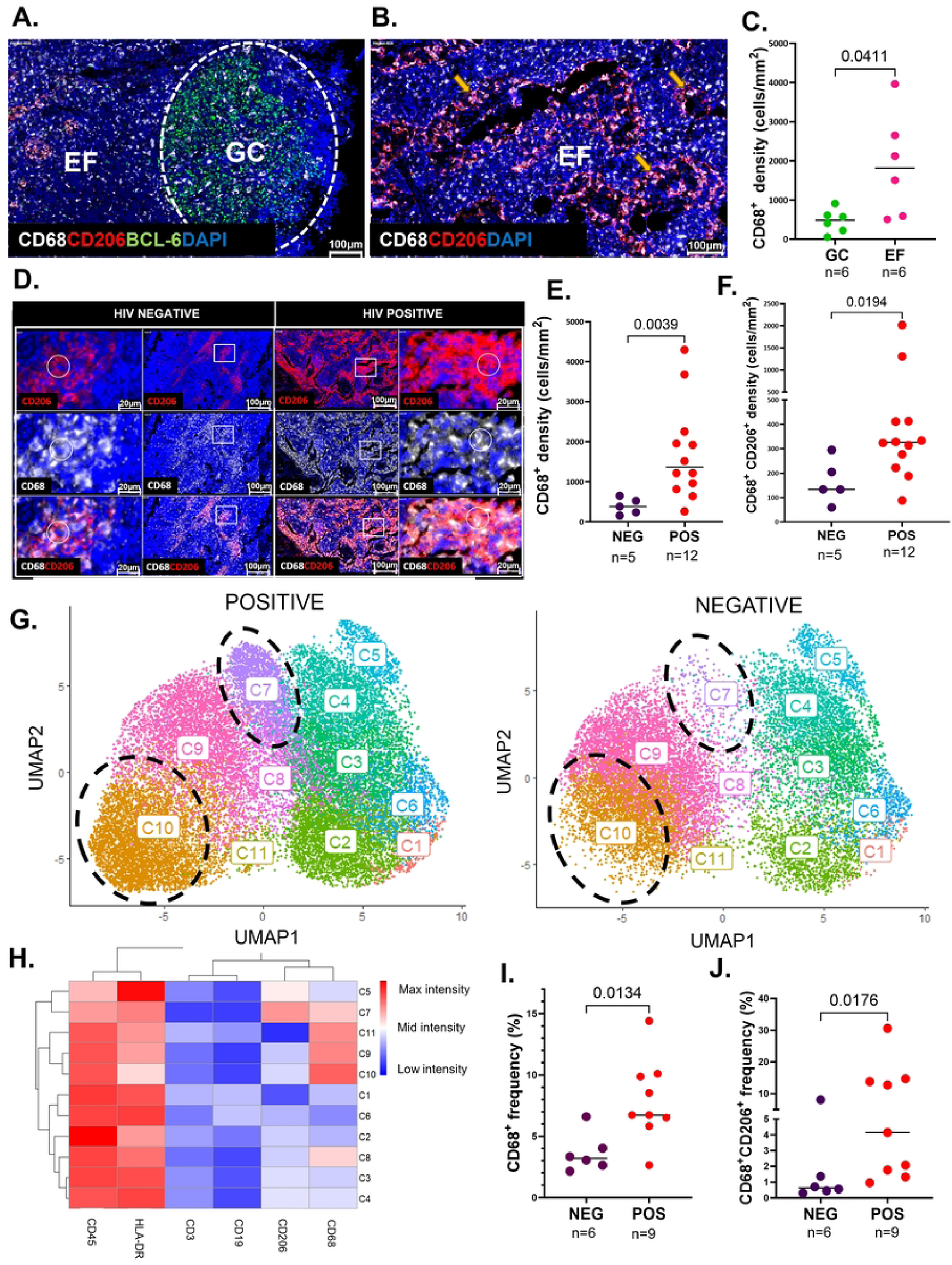
Impact of HIV infection on the phenotypes, location and frequencies of macrophages in human LN tissues. **(A)** Representative multicolor IF image of the localization of CD68^+^ (white) and CD206^+^ (red) macrophages relative to the LN GC (white dashed oval) indicated by BCL-6 (green) positivity or the extrafollicular (EF) region (outside the GC). **(B)** Representative multicolor IF image of CD68^+^CD206^+^ macrophages along LN lymphatic and blood vessels (yellow arrows). **(C)** Aggregate data from quantitative image analysis by TissueQuest (TissueGnostics, Vienna) indicating the density of CD68^+^ macrophages (cells/mm^2^) which localized inside the GC (green) and the EF region (pink) from LN tissues of n=6 different donors, p=0.0411. **(D)** Multicolor IF images of the individual channels of CD206^+^ macrophages (red), CD68^+^ macrophages (white), and the composite image of CD68^+^CD206^+^ macrophages in LN tissue sections from a representative PLWoH and a PLWH. The white rectangles from the inner panels indicate regions of interest which are further magnified on the outer panels. The white circles on the outer panels indicate the double positive CD68^+^CD206^+^ macrophages**. (E-F)** Quantitative image analysis by TissueQuest (TissueGnostics, Vienna) between n=5 PLWoH (purple) and n=12 PLWH (red) of the frequency (cells/mm^2^) of: CD68^+^ macrophages, p=0.0039 **(E)** and CD68^+^CD206^+^ macrophages, p=0.0194 **(F)**. **(G)** Multiparametric global Uniform Manifold Approximation and Projection (UMAP) visualization of LNMCs from 9 PLWH and 6 PLWoH. The Flow Self-Organizing Map (FlowSOM) was overlaid on the UMAP which identified 11 clusters of LNMCs and are labelled on the UMAP. **(H)** The unsupervised hierarchical clustering heatmap summarizes the mean fluorescence intensities of each loaded parameter (CD3, CD19, CD45, HLA-DR, CD68 and CD206), whereby warm colors (dark red) indicate high expression and cold colors (dark blue) indicate low expression. **(I-J)** Frequency of parent (%) comparison between n=6 PLWoH (purple) and n=9 PLWH (red) of CD68^+^ macrophages, p=0.0134 **(I)** and CD68^+^CD206^+^ macrophage frequency, p=0.0176 **(J)** within LNMCs. Images were acquired at 40x and all cell nuclei were detected by DAPI (blue). Scale bars= 100μm and 20μm. Comparisons made using the Mann Whitney tests, whereby horizontal bars denote the median and each circle represents an individual donor. Levels of significance are as follows: *: p<0.05; **: p<0.01 and ***: p<0.001.

Chronic HIV infection is characterized by persistent inflammation, chronic immune activation and T cell dysfunction (31, 32). While the impact of HIV on CD4^+^ T cells is well-documented, the effects on macrophage frequencies and functions in tissues are less understood. In organs such as the lungs and liver, an accumulation of macrophages has been noted even during suppressed HIV infection (33). HIV infection has been shown to initially promote a pro-inflammatory macrophage phenotype, which gradually shifts to an anti-inflammatory phenotype over time (11, 34). However, the specific effects of HIV on macrophage polarization, frequency, and function within LNs remain unclear. We examined the effect of HIV on macrophage polarization and frequency in LN tissues using multicolor IF microscopy combined with TissueQuest quantitative image analysis **(S2 Fig)**. Representative IF staining for CD68, CD206, and DAPI from n=3 PLWoH and n=3 PLWH is shown **(Fig 1D, and S3 Fig)**. We calculated macrophage density (cells/mm²) in 5 PLWoH and 12 PLWH using TissueQuest image analysis software. Aggregate data for 12 PLWH show greater densities of CD68 and CD206 single and double positive populations compared to 5 PLWoH (CD68^+^ p=0.0039, and CD68^+^CD206^+^ p=0.0194) **(Fig 1E-F)**.

While immunofluorescence microscopy is sensitive, it is typically low-plex, and can only detect a few fluorescent markers simultaneously (35). To expand on these findings, we isolated lymph node mononuclear cells (LNMCs) from 9 PLWH and 6 PLWoH and performed detailed macrophage phenotype analysis by flow cytometry. Using the gating strategy illustrated in **(S3 Fig)**, macrophages were defined as CD3^-^CD19^-^CD45^+^HLA-DR^+^ cells (36). From this total macrophage pool, we further classified cells into pro-inflammatory (CD3^-^CD19^-^CD45^+^HLA-DR^+^CD68^+^CD206^-^) and anti-inflammatory (CD3^-^CD19^-^ CD45^+^HLA-DR^+^CD68^+^CD206^+^) subsets. To characterize the diversity of macrophage subsets and examine potential HIV-related changes in their frequencies, we applied UMAP for unsupervised dimensionality reduction and clustering. The UMAP analysis was initially conducted on concatenated LNMC data from the merged study participants n=15 (n=6 PLWoH and n=9 PLWH) to illustrate macrophage sub-population distribution. These populations were clustered with the FlowSOM algorithm which identified 11 distinct populations based on their relative marker expression **(Fig 1G)**. The unsupervised hierarchical clustered heat map categorized the LNMC populations into 11 cell clusters **(Fig 1G-H)**. CD45^+^HLA-DR^+^CD3^-^CD19^-^CD68^+^CD206^-^ pro-inflammatory macrophages were detected in cell clusters C10. CD45^+^HLA-DR^+^CD3^-^CD19^-^CD68^+^CD206^+^ anti-inflammatory macrophages were detected in cell cluster C7 **(Fig 1G)**. Clear distinctions in cell clustering were observed: C7 cells, denoting a CD68^+^CD206^+^ phenotype, and C10 cells, denoting a CD68^+^CD206^-^ phenotype, both showed greater densities in PLWH than in PLWoH (**Fig 1G**). Quantitative analysis confirmed the elevated levels of these two populations in PLWH: CD68^+^CD206^-^ (C10) (p=0.0134) **(Fig 1I)** and CD68^+^CD206^+^ (C7) (p=0.0176) **(Fig 1J)**, supporting the imaging data. Our findings reveal that HIV infection drives significant macrophage expansion in human LNs, which persists even with early ART initiation.

### HIV-1 persists in CD68^+^ lymph node germinal center macrophages despite early ART initiation

The early initiation of ART limits the establishment of the HIV reservoir but does not prevent seeding or eradication (37–39). While it is well established that LNs are a major site of HIV persistence during ART, the precise cellular reservoir phenotypes are poorly characterized. Given the observed increased frequencies of macrophages in LN tissues of PLWH, we sought to characterize the phenotype and topology of macrophage reservoirs in LNs. 12 PLWH were included in these investigations. Selection criteria were based on tissue quality, the presence of GCs, and HIV treatment/suppression status. Multicolor IF microscopy on FFPE LN tissues from these PLWH was performed using the transcription factor BCL-6 to detect active GCs, CD68 to define pro-inflammatory macrophages and Gagp24 to detect HIV protein. CD68^+^ macrophages staining positive for HIV Gagp24 protein were readily detectable, almost exclusively in LN GCs as shown in the representative image from a fully suppressed donor **(Fig 2A)**. 3 LN tissue samples obtained from PLWoH were included as negative controls, and none of them exhibited Gagp24 staining **(S4 Fig)**. We previously demonstrated that HIV primarily persists in LN GCs despite ART initiation in hyperacute infection and plasma viral suppression in PLWH subtype C (5). We also reported reduced total HIV DNA in individuals who initiate therapy during hyperacute HIV infection comparted to those who delay treatment initiation (39). Therefore, we assessed the macrophage reservoir size based on timing of treatment initiation and peripheral virus suppression status. Representative images show that HIV Gagp24^+^ macrophages were present in both acute-treated and chronic-treated participants **(Fig 2B and Fig S4)**. Quantitative analysis by TissueQuest revealed a lower frequency of Gagp24^+^ macrophages in acute-treated donors compared to chronic-treated donors (p=0.0025) **(Fig 2C)**. No significant difference was observed, however, between individuals with complete plasma viral suppression and those who were viremic (p=0.7302) **(Fig 2D)**. Notably, the density of HIV Gagp24^+^ macrophages correlated with pVL (p=0.0008; r=0.9923) **(Fig 2E)**.

**Figure 2.**
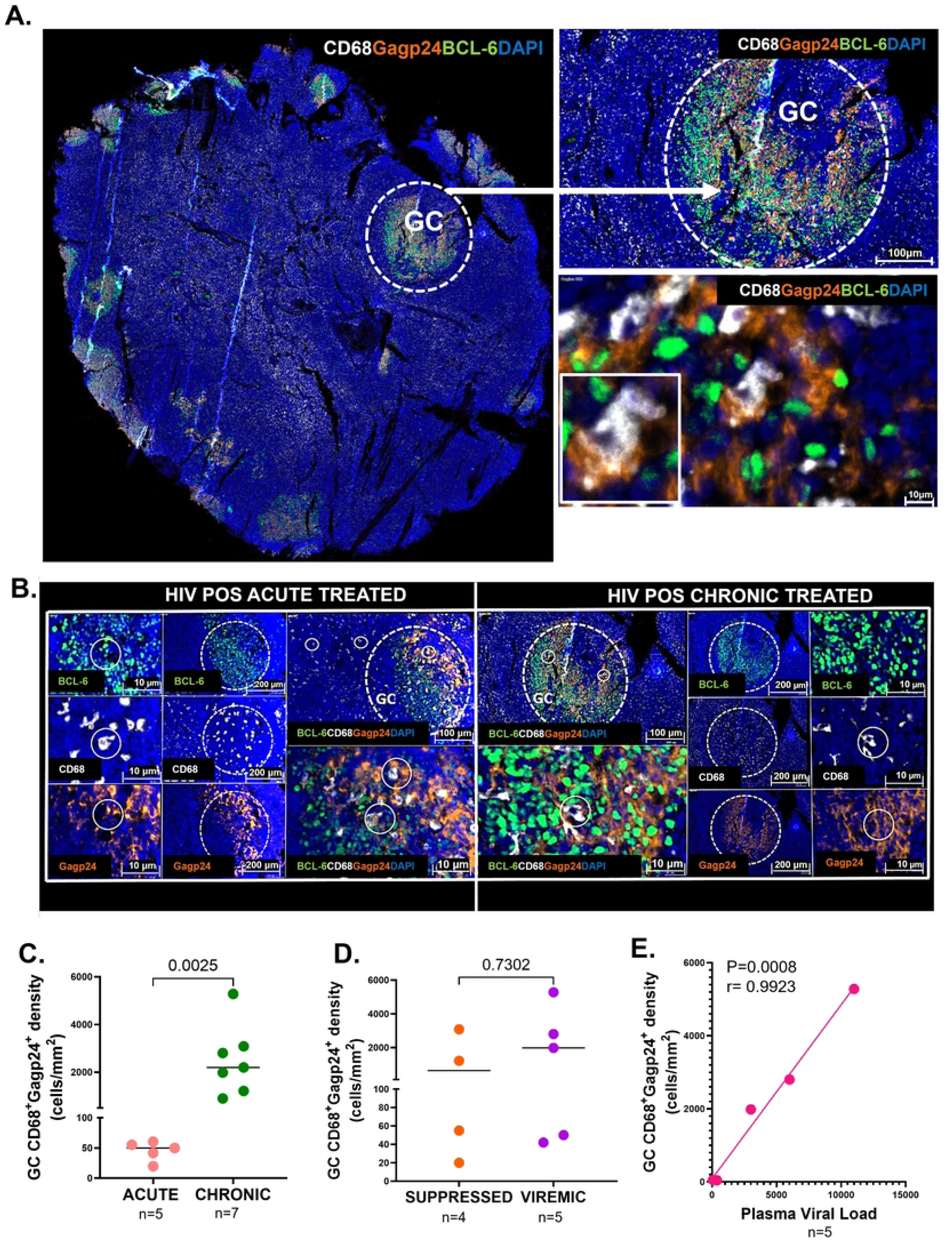
HIV-1 persists in CD68^+^ lymph node germinal center macrophages despite early ART initiation. **(A)** Representative multicolor IF image of a whole LN tissue section of CD68^+^ (white) macrophages, HIV Gagp24 antigen (brown) and BCL-6 (green) to define GCs. A magnified GC (top-right inset) shows co-localization of CD68^+^ macrophages with HIV Gagp24 antigens within the GC, further magnified (bottom-right inset). **(B)** Representative multicolor IF images from an acute-treated and chronic-treated PLWH of GCs (white dashed ovals) indicated by BCL-6 (green) positivity and their respective expressions of CD68^+^ (white) macrophages and HIV Gagp24 antigen (brown). Co-staining of Gag p24 (brown) and CD68 (white) was observed in both acute-treated and chronic-treated PLWH (white solid circles). **(C)** Quantitative image analysis by TissueQuest (TissueGnostics, Vienna) was used to assess and compare the density of GC CD68^+^Gagp24^+^ macrophages (cells/mm^2^) in n=5 acute-treated (peach) and n=7 chronic-treated (green) PLWH, p=0.0025 **(D)** Aggregate quantitative image analysis data by TissueQuest (TissueGnostics, Vienna) comparing the density of GC CD68^+^Gagp24^+^ macrophages (cells/mm^2^) in n=4 virally suppressed (orange) and n=5 viremic (purple) PLWH, p=0.7302. **(E)** Correlation between the density of GC CD68^+^Gagp24^+^ macrophages (cells/mm^2^) and plasma viral load (VL) in n=5 PLWH (pink), p=0.0008, r=0.9923 by Spearman’s ranked correlation analysis. Images were acquired at 40x magnification and all cell nuclei were detected by DAPI (blue). Scale bars equal to 10μm, 100μm and 200μm. Comparisons made using the Mann Whitney tests, whereby horizontal bars denote the median and each circle represents an individual donor. Levels of significance are as follows: *: p<0.05; **: p<0.01 and ***: p<0.001.

### Lymph node tissues harbor transcriptionally active macrophage reservoirs primarily in BCFs

Building on the observed co-localization of LN GC CD68^+^ macrophages with HIV Gagp24, we analyzed their capacity to harbor transcriptionally active HIV reservoirs. We employed a fluorescence *in situ* hybridization (FISH) protocol (RNAscope), along with multicolor IF microscopy (5, 29, 40, 41), to identify and quantify macrophages harboring transcriptionally active HIV reservoirs. Initial validation of our staining was performed on LN tissues using positive and negative control RNA probes, which yielded expected results **(Fig 3A)**. Based on prior staining with Gagp24, treatment status and sample availability, we stained LN tissues from 5 participants of whom 20% were virally suppressed and 80% were viremic. Consistent with Gagp24 protein staining, the HIV-1 *gag-pol* RNA (vRNA) signal was predominantly localized in the GCs **(Fig 3B)**. Representative images from two donors **(Fig 3C)** and supplementary images from three additional donors **(S5 Fig)** demonstrate that CD68^+^ macrophages containing vRNA were identified in 4/5 samples analyzed. We next utilized the FISH-IF module within HALO image analysis (version v3.6.4134.396; Indica Labs) to assess the contribution of macrophages to transcriptionally active reservoirs within GCs. Aggregated data from 8 GCs across 4 PLWH revealed that, on average, 32.93% of GC-resident cells harbor vRNA, with 8.99% of these cells being CD68^+^ macrophages **(Fig 3D)**. Among the total CD68^+^ macrophage population in the GC a median of 44.05% harbor vRNA **(Fig 3E)**. Individual RNAscope H-scores determined by HALO are illustrated **(S5 Fig)**.

**Figure 3.**
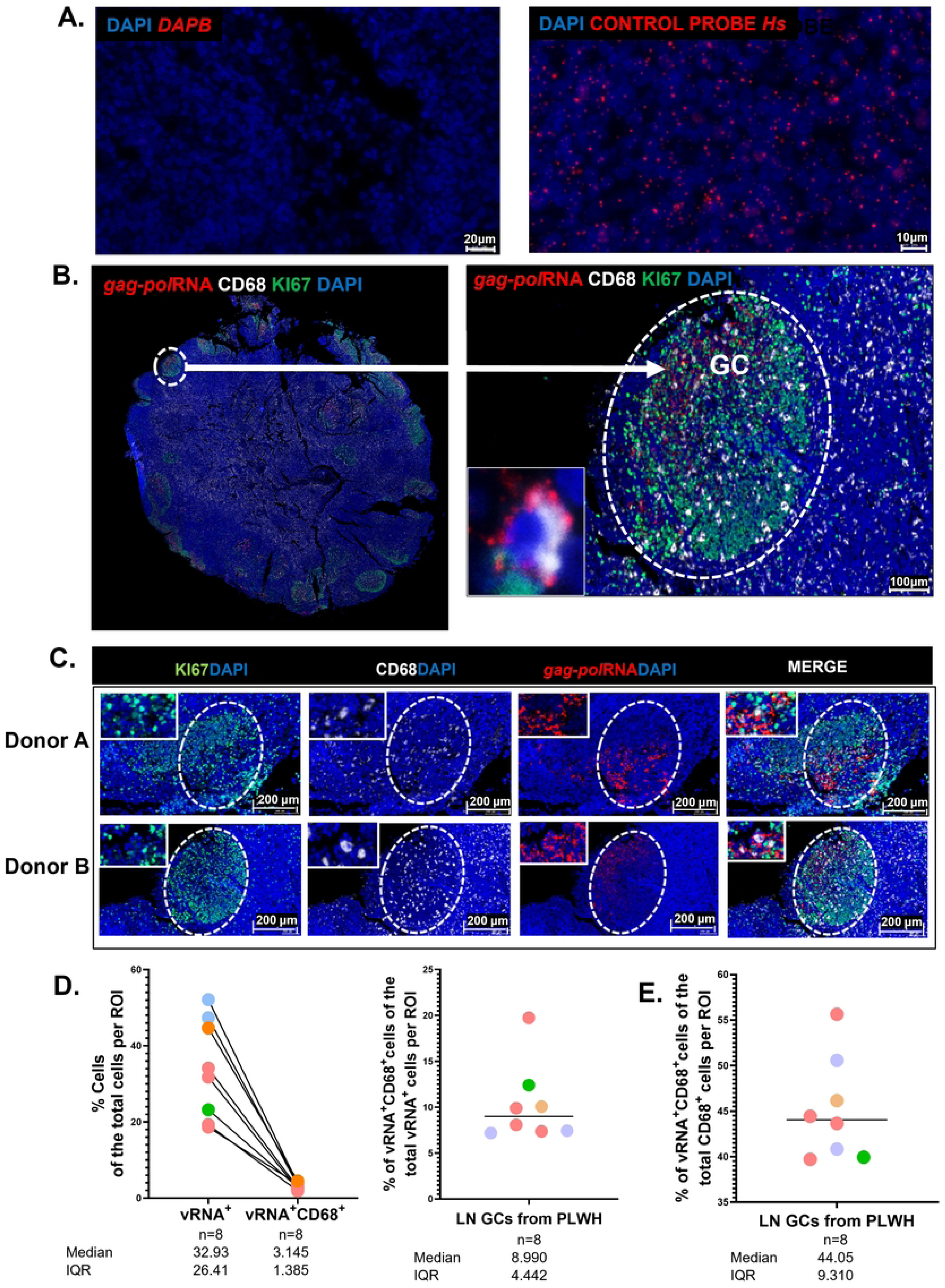
Lymph node tissues harbor transcriptionally active macrophages reservoirs primarily in BCFs. **(A)** Fluorescence in situ hybridization (FISH) RNAscope negative and positive control images using the *DAPB* (red) or Control Probe *Hs* (red) respectively. **(B)** Representative FISH RNAscope coupled with multicolor IF image of a whole LN tissue section of CD68^+^ (white) macrophages, HIV *gag-pol* RNA (red) and Ki67 (green) to define GCs (white dashed ovals). A magnified GC (right inset) shows co-localization of CD68^+^ macrophages with HIV *gag-pol* RNA within the GC, further magnified (bottom left inset). A CD68*^+^* macrophage (white) is seen co-localizing with HIV *gag-pol* RNA transcripts (red). **(C)** Representative images of FISH RNAscope coupled with multicolor IF of *gag-pol* HIV-1 RNA (red), KI67 (green), and CD68 (white) detection within LN GCs in n=2 PLWH showing the individual channels and composite image. The white dashed oval defines the GC. Single RNA transcripts are shown as punctate dots. **(D)** Aggregate data from HALO image analysis quantification of LN tissues from n=8 GCs from n=4 PLWH of the %*gag-pol*RNA^+^ (vRNA^+^) and %vRNA^+^CD68^+^ cells out of the total cells per region of interest (ROI). The proportion of vRNA^+^CD68^+^ cells out of the total vRNA^+^ cells per ROI is shown alongside. **(E)** Aggregate data from HALO image analysis quantification of LN tissues from n=8 GCs from n=4 PLWH of the percentage of vRNA^+^CD68^+^ cells out of the total CD68^+^ cells per ROI is shown. Images acquired at 40x and all cell nuclei were detected by DAPI (blue). Scale bars= 10μm, 20μm, 100μm and 200μm. Each colored circle from the aggregate data represents an individual LN tissue donor. Multiple dots of the same color represent multiple regions of *gag-pol* RNA positivity from the same donor. Levels of significance are as follows: *: p<0.05; **: p<0.01 and ***: p<0.001.

### Detection of HIV subtype C proviral DNA within lymph node macrophages

While both HIV Gagp24 protein and vRNA detection identify HIV infected macrophages, these findings do not directly indicate the presence of latent macrophage proviral reservoirs, which are characterized by integrated HIV DNA (1). To better define proviral macrophage reservoirs, we first used the highly sensitive *in situ* FISH DNAscope assay (40, 41). Here, we employed an HIV *gag-pol* sense probe optimized for HIV Clade C, which had a 92.8% alignment with sequences from our study cohort **(S5 Fig)**. We validated the specificity of the DNA probe using positive and negative control cell lines **(Fig 4A)**. The analysis included 6 LNs from PLWH: 3 from virally suppressed and 3 from viremic participants who were selected based on peripheral suppression status, detectable GCs and recent excisional dates, as well as 3 PLWoH. As expected, no vDNA was detected in the HIV-negative LNs **(Fig 4B and S5 Fig)**. Representative images **(Fig 4C)** and aggregate data **(Fig 4D)** show that all 6 LNs from PLWH contained varying levels of detectable subtype C gag-pol DNA (vDNA). Proviral DNA punctate signals were predominantly localized within the nuclei of CD68^+^ macrophages from PLWH **(Fig 4C)**. Next, we wanted to investigate the impact of spatial localization on the contribution of macrophages to the proviral reservoir in hotspot vDNA^+^ zones within LN tissues. Here, we used the FISH-IF module in HALO to segment and analyze 1-3 GC ROIs (based on the frequency of GCs per LN tissue) and 3 EF ROIs with an average cell count of 1500 cells per ROI in PLWH. Spatial analysis revealed that, unlike viral RNA and protein, the DNA signal was found in both follicular and EF zones, where the median proportion of vDNA^+^ macrophages of the total vDNA^+^ cells per zone was as follows: GC=0.077 and EF=0.150 (p=0.0074) **(Fig 4E)**. We then assessed the impact of plasma suppression status on the proportion of vDNA^+^ macrophages of the total vDNA^+^ cells per zone and found no significant difference (p=0.3508) **(Fig 4F)**. Additionally, we observed multiple punctate vDNA signals within the nuclei, which are thought to represent different HIV integration sites across the genome (42). With the rationale that viremic PLWH have a greater probability of viral re-activation events, we compared vDNA copies per cell in the viremic and virally suppressed groups and observed no significant difference (p=0.9922) **(Fig 4G)**.

**Figure 4.**
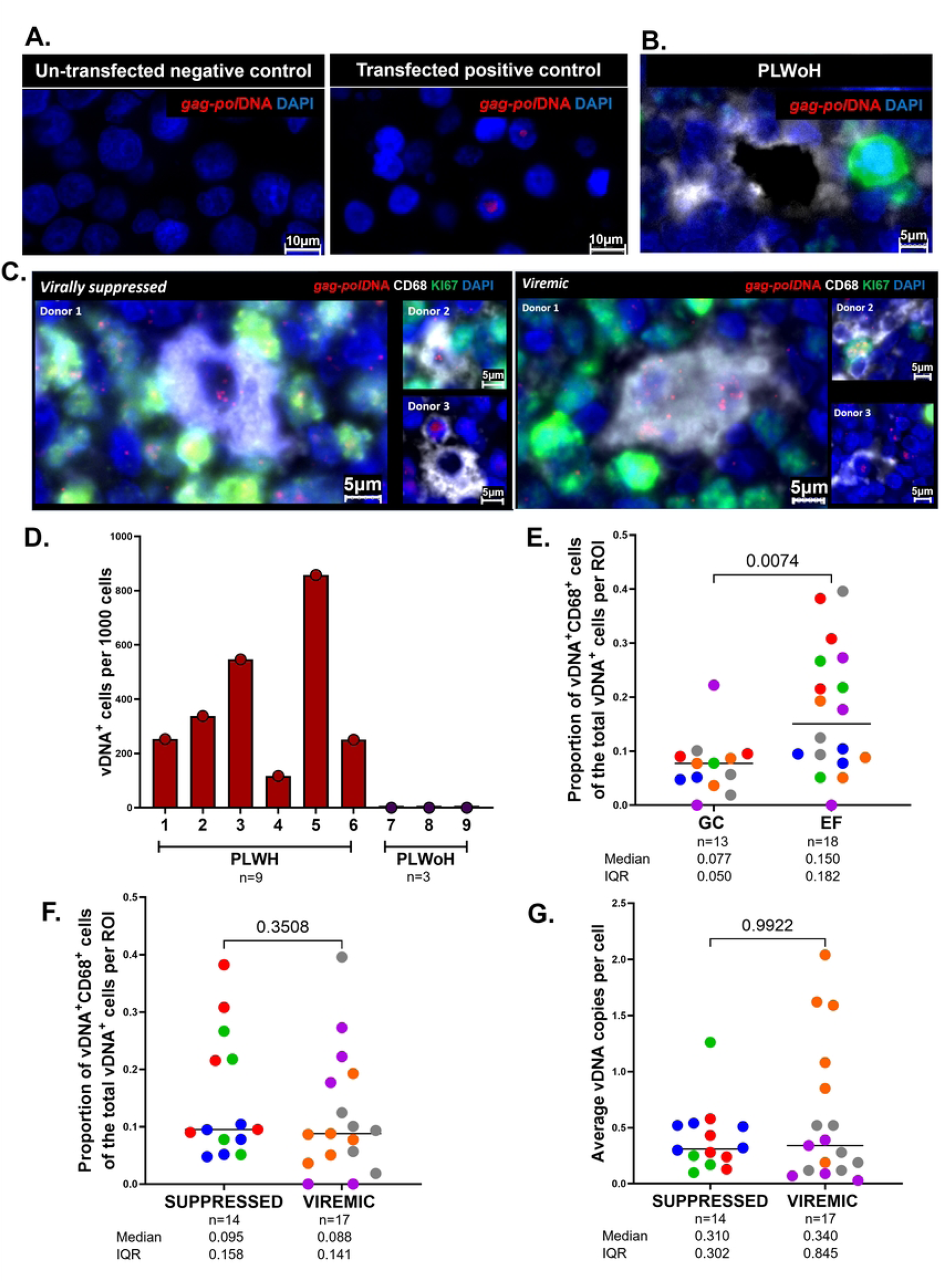
Detection of HIV subtype C proviral DNA within lymph node macrophages. **(A)** FISH DNAscope of *gag-pol* HIV DNA (red) in negative and positive controls using un-transfected and HIV-transfected cell lines respectively. **(B)** Representative FISH DNAscope image from a PLWoH of *gag-pol* HIV DNA (red), CD68 (white), KI67 (green) and nuclei counterstained in DAPI (blue). **(C)** Representative images of FISH DNAscope coupled with multicolor IF of *gag-pol* HIV subtype C DNA (red), Ki67 (green), and CD68 (white) detection within LN tissues from n=3 virally suppressed and n=3 viremic PLWH. **(D)** Aggregate data of the frequency of *gag-pol*DNA^+^ (vDNA^+^) cells per 1000 cells between n=6 PLWH (red) and n=3 PLWoH (purple) using HALO image analysis. **(E)** The proportion of vDNA^+^CD68^+^ cells out of the total vDNA^+^ cells per ROI between n=13 GC and n=18 EF regions from n=3 suppressed and n=3 viremic PLWH, p=0.0074. **(F)** The proportion of vDNA^+^CD68^+^ cells out of the total vDNA^+^ cells per ROI between n=14 regions from n=3 suppressed and n=17 regions from n=3 viremic PLWH, p=0.3508. **(G)** Average vDNA copies per cell between n=14 regions from n=3 suppressed and n=17 regions from n=3 viremic PLWH, p=0.9922. Images were acquired at 40x and all cell nuclei were detected by DAPI (blue). Scale bars= 5μm and 10μm. Comparisons made using the Mann Whitney tests, whereby horizontal bars denote the median and each color represents an individual donor and circle represents an individual ROI. Levels of significance are as follows: *: p<0.05; **: p<0.01 and ***: p<0.001.

### HIV proviral DNA quantification and comparison between lymph node myeloid and CD4^+^ T cells

To further evaluate the potential role of macrophages as latent reservoirs, we measured total proviral DNA levels in myeloid cells and CD4^+^ T cells derived from LNs using ddPCR, which is typically used to measure total proviral reservoir (39). CD4^+^ T cells and paired myeloid cell subsets were FACS-sorted from LNMCs, as outlined in the gating strategy **(S6 Fig)**. We sorted the total myeloid population rather than macrophage subsets due to the insufficient yield of purified macrophage populations from the tissues required for the ddPCR assay (7). 10 PLWH comprising of 6 suppressed and 4 viremic participants were selected based on peripheral suppression status and sample availability and used for these studies. Next, we conducted the ddPCR assay to detect and quantify HIV proviral DNA levels (log copies per million cells) within these cellular compartments (39) as shown by the representative ddPCR 1-D plot from a viremic PLWH **(Fig 5A)**. Assay validatory checks by TREC PCR revealed that there was no CD4^+^ T cell contamination within the sorted myeloid cells **(Fig 5B)**. Assay control testing from n=4 PLWoH yielded no detectable HIV DNA as expected **(Fig 5C)**. We detected proviral HIV DNA in the CD4^+^ T cell population in all 10 PLWH. In the myeloid compartment, proviral HIV DNA was detected in 7 out of 10 PLWH (3/6 virally suppressed and 4/4 viremic PLWH). In virally suppressed PLWH (pVL<20 copies/mL) the proportion of HIV DNA levels was higher in the CD4^+^ T cell population (98.4%) compared to the myeloid cell population (1.6%) (p=0.0532) **(Fig 5D)**. In viremic PLWH (pVL>250), we observed a similar trend of higher HIV DNA levels in the CD4^+^ T cell population (71.3%) compared to the myeloid cell population (28.7%) (p=0.0940) **(Fig 5E)**. Notably, the proportion of proviral DNA from the myeloid compartment was higher in viremic PLWH compared to suppressed PLWH **(Fig 5D-E)**. HIV DNA levels were similar in viremic compared to suppressed PLWH in the pooled CD4^+^ T cell and myeloid cell population (p=0.2665) **(Fig 5F)** and within the CD4^+^ T cell population (p=0.1939) **(Fig 5G)**. However, in the myeloid cell population, HIV DNA levels were higher in viremic compared to virally suppressed PLWH (p=0.0190) **(Fig 5H)**. Our findings reveal that the myeloid compartment harbors about 29% of the total proviral DNA in LNs of viremic individuals. In contrast, the myeloid compartment harbors only 1.6% of the total proviral DNA in LNs of virally suppressed PLWH, suggesting a marked reduction in proviral burden within myeloid cells during suppression.

**Figure 5.**
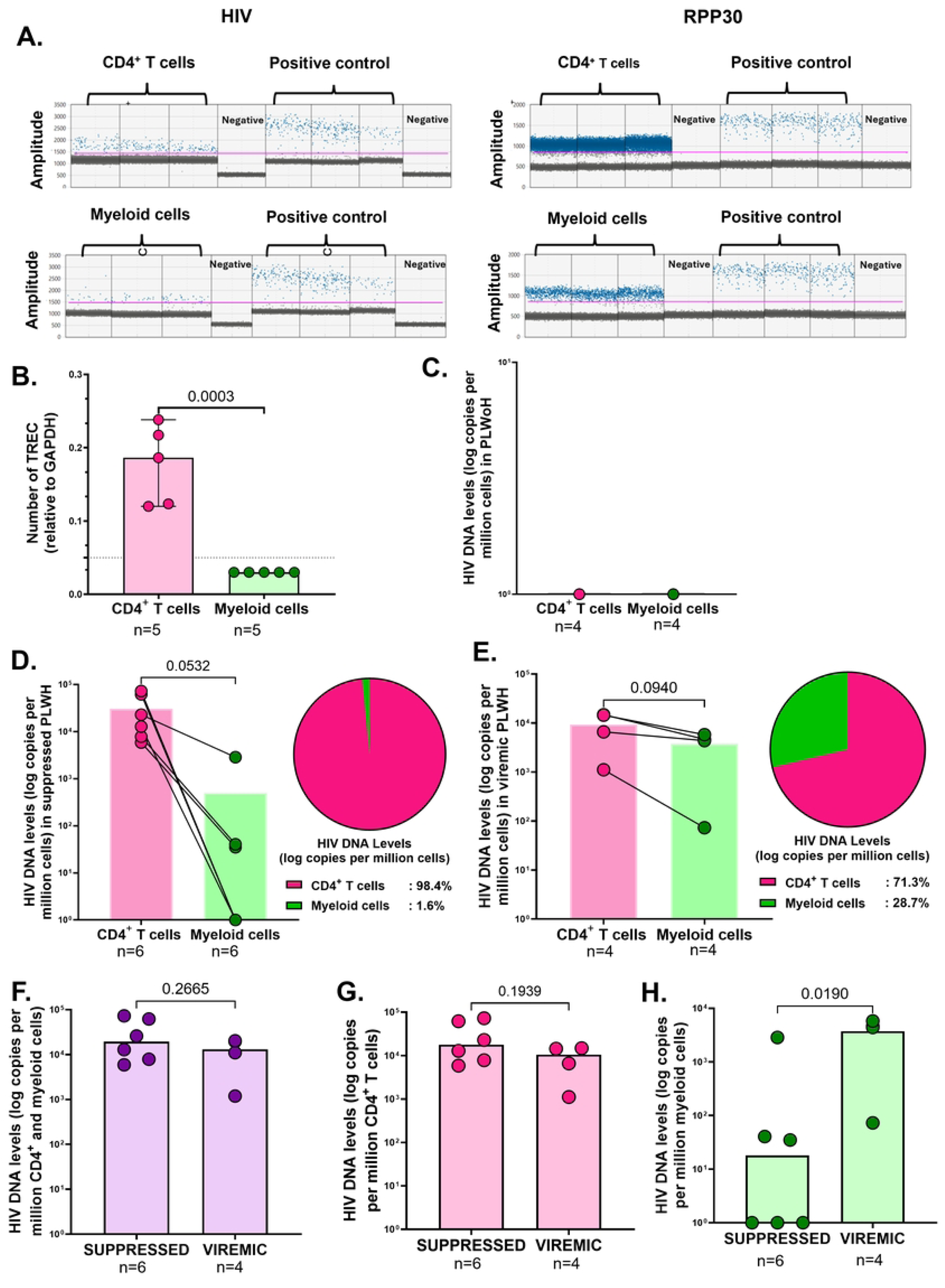
HIV proviral DNA quantification and comparison between lymph node myeloid and CD4^+^ T cells. **(A)** Representative 1-D ddPCR plot from a viremic PLWH showing droplets from CD4^+^ T and myeloid cell samples, with each droplet from a sample plotted on a graph of fluorescence intensity versus droplet number. Blue dots denote positively amplified droplets while grey dots are negative. A manual threshold indicated by the pink line was set to distinguish the positive and negative droplet clouds. DNA from the 8E5/LAV cell line was used as a positive control. **(B-C)** ddPCR controls: Number of TREC relative to GAPDH **(B)** and HIV DNA levels (log copies per million cells) in n=4 PLWoH between CD4^+^ T cells (pink) and myeloid cells (green) **(C)**. **(D)** Aggregate data comparing HIV DNA levels (log copies per million cells) in n=6 virally suppressed PLWH between CD4^+^ T cells (pink) and myeloid cells (green), p=0.0532 by Student’s paired t test. The pie chart (right) depicts the proportion of HIV DNA levels in the CD4^+^ T cells (pink) compared to myeloid cells (green) in virally suppressed PLWH. **(E)** Aggregate data comparing HIV DNA levels (log copies per million cells) in n=4 viremic PLWH between CD4^+^ T cells (pink) and myeloid cells (green), p=0.0940 by Student’s paired t test. The pie chart (right) depicts the proportion of HIV DNA levels in the CD4^+^ T cells (pink) compared to myeloid cells (green) in viremic PLWH. **(F-H)** Aggregate data comparing HIV DNA levels (log copies per million cells) in n=6 virally suppressed PLWH and n=4 viremic PLWH, in pooled CD4^+^ T cells and myeloid cells, p=0.2665 by Students unpaired t test (F), in isolated CD4^+^ T cells, p=0.1939 by Students unpaired t test **(G)** and in isolated myeloid cells p=0.0190 by Mann Whitney test. **(H)** Levels of significance are as follows: *: p<0.05; **: p<0.01 and ***: p<0.001.

### Distinct spatial distribution and morphology of CD68^+^ macrophages exhibiting CD4^+^ T cell phagocytosis and HIV productive infection

Although evidence supporting the existence of macrophage reservoirs is growing, the significance of these reservoirs in HIV persistence remains unsettled (43). A primary point of contention is accurately distinguishing macrophages harboring productive HIV infection from those that have merely phagocytosed HIV-infected CD4^+^ T cells (44, 45). To address this issue, we used high resolution IF microscopy (100x oil immersion objective) to visualize CD4^+^ T cell phagocytosis by CD68^+^ macrophages and distinguish these from productively infected macrophages without evidence of CD4^+^ T cell ingestion. We selected LN tissues from 11 PLWH based on sample availability, presence of GCs and treatment status. Of these participants, 5 were acute-treated and 6 were chronic-treated, and 63.64% were suppressed 36.36% were viremic. Intriguingly, we documented multiple events showing the sequential stages of CD4^+^ T cell engulfment and subsequent ingestion by macrophages **(Fig 6A-D)**, starting with CD4^+^ T cell attachment to macrophage membrane during the initial stages of phagocytosis **(Fig 6A)**, followed by partial CD4^+^ T cell engulfment **(Fig 6B)**. Thereafter, complete CD4^+^ T cell ingestion was observed with complete intracellular CD4 positivity within the CD68^+^ macrophage **(Fig 6C)**. Lastly, aggregation of macrophages that have ingested CD4^+^ T cells occurs **(Fig 6D)**. The CD4^+^T cell ingestion events by macrophages were mostly observed outside the GCs. In contrast, macrophages within GCs stained positive for HIV Gagp24 in the absence of intracellular CD4^+^ T cell staining suggesting productive infection other than CD4^+^ T cell ingestion **(Fig 6E-H)**. All productively infected macrophages were detected in LN GCs and exhibited multinucleated giant cell morphology **(Fig. 6E-H)**. Overall, CD4-ingested macrophages were highly distinguishable from productively infected macrophages in terms of size, morphology, spatial localization relative to LN GCs, CD4 membranous positivity and intracellular CD4 positivity. Our quantitative analysis demonstrated a markedly higher frequency of productively infected macrophages relative to macrophages that had phagocytosed CD4^+^ T cells (p < 0.0001) **(Fig 6I and S6 Fig)**.

**Figure 6.**
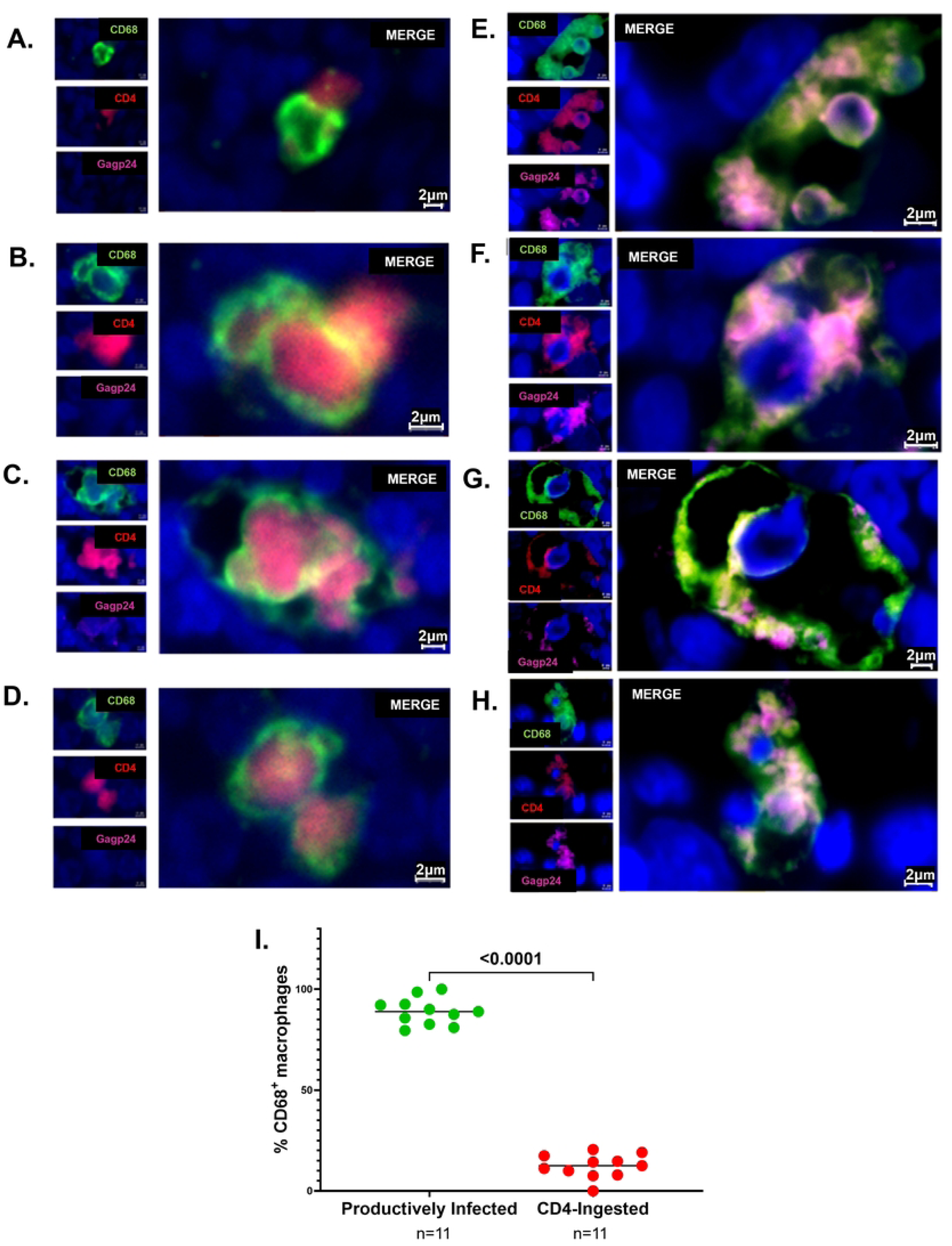
Distinct spatial distribution and morphology of CD68^+^ macrophages exhibiting CD4^+^ T cell phagocytosis and HIV productive infection. Representative multicolor IF high-resolution images of CD68 (green), CD4 (red) and Gagp24 (pink). **(A-D)** The different stages of CD4^+^ T cell ingestion by CD68^+^ macrophages were captured outside the GCs of LN tissues obtained from PLWH on suppressive ART. **(A)** A CD4^+^ T cell (red) attaches to the membrane of the CD68^+^ macrophage (green) indicating phagocytosis initiation. **(B)** Partial CD4^+^ T cell ingestion by the CD68^+^ macrophage – note that intracellular CD4^+^ (red) staining starts to appear within the CD68^+^ (green) macrophage. **(C)** Complete CD4^+^ T cell ingestion by the CD68^+^ cell, as illustrated by the outer membranous staining of CD68 (green) and complete intracellular staining of CD4 (red). **(D)** Cell-cell contact of two ingested CD4^+^ T cells by CD68^+^ macrophages were also captured. **(E-H)** Productively HIV infected CD68^+^ macrophages are multinucleated giant cells which co-localize with HIV Gagp24 in the absence of intracellular CD4 positivity. **(I)** Percentage of productively HIV infected CD68^+^ macrophages (green) and CD4^+^ T cell-ingested CD68^+^ macrophages (red) from n=11 LN tissue samples from PLWH on suppressive ART, p<0.0001 by Mann-Whitney test. Each circle represents an individual LN tissue donor and the horizonal bars denote the median. Images were acquired at 100x using oil immersion and all cell nuclei were detected by DAPI (blue). Scale bars= 2µm. Levels of significance are as follows: *: p<0.05; **: p<0.01, ***: p<0.001 and ****: p<0.0001.

## Discussion

The eradication of HIV remains challenging due to the virus’s ability to establish and persist in reservoir sites, particularly within lymphoid tissues (1, 5). Understanding the role of lesser-studied cell types, such as macrophages, in maintaining HIV reservoirs is essential for achieving complete HIV eradication. In this study, we characterized the phenotype, localization, and reservoir nature of macrophages in the LN tissues of individuals with suppressed HIV clade C infection (1, 46). Our findings demonstrate that LN macrophages harbor both transcriptionally active and proviral reservoirs, highlighting their critical role in sustaining tissue-based viral reservoirs during ART. Importantly, we reveal significant contributions of macrophages to the LN proviral and active reservoirs through robust *in situ* quantitative analysis.

Quantitative ddPCR analysis revealed that when compared to CD4^+^ T cells, myeloid cells account for approximately 1.6% of the proviral HIV reservoir in LNs of virally suppressed individuals and approximately 28.7% in viremic PLWH. Previous findings for urethral macrophages show that around 1% of macrophages contain integrated proviral DNA and 0.2% to 0.9% of macrophages contain active forms of the reservoir (7). In this study we show by quantitative image analysis that vRNA^+^ macrophages account for approximately 3.14% of cells within the follicle, whereby a median of 44.05% of follicular macrophages contain vRNA transcripts. These results are consistent with previous data showing that CD11c^+^CD68^+^ cells represent about 7% of total cells in the GC (47). While, the data shows that the latent and active reservoir size differs across multiple tissue sites, with a greater proportion harbored by LNs (41, 48), it is important to note that current image-based measurements of reservoirs likely overestimate the true replication-competent reservoirs capable of reactivation after treatment interruption. This underscores the necessity of developing advanced assays to distinguish defective proviral DNA from intact sequences *in situ* with greater precision (49, 50).

Although the myeloid reservoir makes up a smaller portion of the overall HIV reservoir compared to CD4^+^ T cells (20), individuals who delay ART initiation show a significant increase in the ratio of HIV DNA in myeloid cells relative to CD4^+^ T cells. We recently reported that whilst acute-treated PLWH had a steady decline in total HIV DNA, total HIV DNA remained stable in chronic-treated PLWH due to slower decay of intact and defective viral genomes compared to acute-treated PLWH (51). This suggests that delayed initiation of ART favors an increased contribution of macrophages to the overall HIV reservoirs in tissues.

Spatial analysis revealed that proviral macrophage reservoirs can be found in both follicular and EF sites, indicating an absence of immune pressure on these silent reservoirs (52). Recently, Liu et al., spatially resolved the whole transcriptomic profile of CD68^+^ macrophages in reactive lymphoid tissues and discovered distinct functional features of macrophages based on their spatial localization (53). For instance, they found that follicular macrophages upregulate genes that promote cell proliferation and metabolism-associated pathways. In contrast, EF macrophages are enriched with genes that promote IFN-γ responses and TNF-α/NF-κB pathways (53). Notably, higher IFN-γ expression and increased effector-target cell contact time hinders efficient killing of macrophages by CTLs (22), providing insight into the potential mechanism favoring proviral reservoir survival in macrophages within EF regions.

In contrast to transcriptionally silent reservoirs, active macrophage reservoirs in LNs are preferentially localized within follicular regions. This observation is consistent with earlier studies indicating that HIV RNA predominantly accumulates in LN GCs (5, 21, 42). This preferential localization is thought to result from CTL-driven clearance of infected cells in EF regions (6, 22, 54, 55). Overall, these observations imply that immune surveillance preferentially eliminates transcriptionally active reservoirs in EF regions, confining them to immune-privileged GCs, whereas, transcriptionally silent reservoirs persist as stable reservoirs (52).

Macrophages are generally less vulnerable to infection by CCR5-utilizing T-tropic HIV, which is the primary form of transmitted and circulating viruses (56). However, studies indicate that macrophages can be infected early by T-tropic viruses, primarily through the phagocytosis of infected CD4^+^ T cells, whereby some virions escape lysosomal degradation, resulting in productive infection (44, 56). Macrophages can also be directly infected by M-tropic viruses (45, 57–59), but, a clear distinction between productive infection and ingestion of HIV-infected CD4^+^ T cells has not been previously demonstrated through imaging techniques. Moreover, *in vitro* infection assays used to classify the HIV-1 cellular M-tropism of the viral isolates do not fully recapitulate the different modes of macrophage infection *in vivo* (11, 58).

This study used high-resolution microscopy to visually distinguish CD4^+^ T cell ingestion by macrophages from productive macrophage infection *in vivo*. By tracking the sequence of events associated with CD4^+^ T cell uptake, we were able to discern these processes based on distinct morphological, phenotypic, and spatial characteristics. Consistent with previous findings, productively infected follicular macrophages were larger compared to CD4-ingested cells (8, 60, 61). These giant multinucleated macrophages have been reported in the literature and are referred to as ‘tingible body macrophages’ (TBMs) (8, 60, 61). They reside in GCs where they capture and clear apoptotic B cells (61–63). The mechanism by which these CD68^+^ TBMs expand requires further interrogation using specific apoptotic markers. Additional mechanisms of macrophage infection apart from CD4-ingestion should be investigated in the future such as tunnelling nanotube (TNT) formation (64–69), cell-cell fusion, forming multinucleated giant cells (70–73), as well as *trans*-fection using virus-containing compartments (VCCs) (74–77).

Overall, this report adds to the mounting evidence that macrophages within lymphoid tissues play a pivotal role as reservoirs for HIV. These findings underscore the importance of targeting macrophages in future HIV eradication strategies. However, targeting macrophage reservoirs for elimination will require a multifaceted, tissue-specific approach. For instance, the efficacy of reverse-transcriptase (RT) inhibitors and latency-reversing agents (LRAs) differs between CD4^+^ T cells and macrophages (78, 79). On a positive note, the anti-cancer agent Imatinib was recently shown to inhibit M-CSF receptor activation, restoring the apoptotic sensitivity of HIV-1 infected macrophages whilst sparing uninfected macrophages, suggesting it as a viable option to specifically target macrophage reservoirs (80).

### Limitations of the study

This study employed CD68 and CD206 as markers to characterize macrophage phenotypes and their spatial distribution. While these markers are well-established, it is important to acknowledge that many other markers, such as IL-1R, CD163, and IL-4R, can also be used to define macrophage subsets with greater precision. Notably, CD68, despite being a canonical macrophage marker, can also be expressed by other immune cell types, including certain lymphocytes. To address this limitation, we incorporated morphological analysis and negative CD3 staining to ensure that the cells analyzed were indeed macrophages (7, 47). Additionally, we utilized a second macrophage-specific marker Iba-1 to further validate that the CD68^+^ and CD206^+^ cells studied were bona fide macrophages (7, 81). Although, the validations measures strengthen the reliability of our findings, future studies should incorporate a broader range of macrophage-specific markers to more comprehensively characterize macrophage reservoirs subsets within human LNs. Furthermore, our study was predominantly conducted in women (5/59 participants were men) from KwaZulu-Natal, South Africa, which limits the generalizability of our findings to other populations, particularly men. Future research should include male participants and broader populations to better understand the interplay between sex, genetic variability, and different HIV subtypes on tissue-based HIV reservoir dynamics. Lastly, the difficulty in sorting sufficient tissue macrophages for the measurement of total proviral DNA by ddPCR necessitated the use of myeloid cells in these studies.

## Materials and methods

### Study participant details

All study participants provided written informed consent prior to inclusion in the study. Ethical approval for the study was granted by the University of KwaZulu-Natal Biomedical Research Ethics Committee (protocol number BF298/14) and the Institutional Review Board of Massachusetts General Hospital (protocol number 2015-P001018). The participants included in this study (**Table 1 and S1 Table**) were recruited through two distinct arms. The first arm employed opportunistic recruitment, targeting patients already scheduled for LN excision or endoscopic examination at Prince Mshiyeni Hospital and Durdoc Hospital for diagnostic or therapeutic purposes. The second arm involved recruiting from existing prospective study cohorts within the HIV Pathogenesis Programme (HPP), specifically the Females Rising through Education, Support and Health (FRESH) and ACUTE study cohorts (82). A total of n=59 study participants were grouped as n=45 PLWH and n=14 PLWoH. The PLWH were further stratified into n=10 acute-treated, n=31 chronic-treated. Four PLWH had an unknown treatment status.

## Method details

### Sample Collection

Excisional LN tissues along with paired blood samples, were collected from these participants exclusively for research purposes as part of the Lymph Node Study (83). Selection criteria were applied in selecting participants to ensure their eligibility for the study. The procedures were carried out safely and in accordance with ethical guidelines by surgical teams at both Prince Mshiyeni and Durdoc Hospitals (83). Lymph node tissue samples underwent processing and embedding at our laboratories in the Africa Health Research Institute (AHRI), while collected blood samples were forwarded to Neuberg Global for viral load and CD4 count testing.

### Immunofluorescence (IF) microscopy

Formalin-fixed tissues were first prepared into paraffin-embedded tissue blocks. These FFPE blocks were then sectioned into 4µm slices and affixed onto Surgipath X-tra adhesive pre-cleaned micro slides in preparation for antibody staining. The prepared slides underwent an overnight baking process to soften the paraffin wax and improve tissue adherence onto the glass slide.

Following baking, the slides were deparaffinized in two changes of xylene for 5 minutes each to expose the tissue. Subsequently, the tissue underwent gradual rehydration by immersion in 100% ethanol for 2 minutes, 95% ethanol for 2 minutes, and finally 70% ethanol for 1 minute. Upon rehydration, the tissue was boiled in 1x EnVision FLEX TRS High pH solution for 20 minutes to expose protein epitopes. Endogenous peroxidases were then blocked using REAL Peroxidase-Blocking Solution for 10 minutes, followed by Bloxall® Blocking solution for an additional 10 minutes.

Primary antibody was applied, followed by incubation with Opal polymer HRP Ms + Rb secondary antibody for 20 minutes. Detection was done using the Opal polymer range 520, 570, and 690, 10 mins respectively, for each round of staining, followed by counterstaining with a prepared DAPI solution for 5 minutes. For multiplexing with additional antibodies, the tissue underwent boiling in AR6 buffer for 20 minutes to remove the previous antibody complex while preserving the covalently bound TSA fluorescent label, after which the steps were repeated.

Primary antibodies utilized included anti-human BCL-6 (clone PG-B6p, Dako/Agilent Technologies), Ki67 (IR62661, Dako/Agilant Technologies), CD206 (clone 685645, R&D Systems), CD68 (clone KP1, Dako/Agilent Technologies), and p24 (clone Kal-1, Dako/Agilent Technologies Cell Sciences or clone MAP1341, Abnova).

Imaging was conducted using a Zeiss Axio Observer inverted microscope at 40x magnification equipped with a Hamamatsu C13440-20C camera. This was facilitated by TissueFAXS imaging software from TissueGnostics (Vienna, Austria). Quantitative image analysis was performed using TissueQuest (TissueGnostics). Our lab upgraded the Zeiss Axio Observer inverted microscope to the Zeiss Imager Z.2 microscope (TissueGnostics, Vienna, Austria), and the TissueQuest image analysis software (TissueGnostics, Vienna, Austria) to StrataQuest Image Analysis software (TissueGnostics, Vienna, Austria). These upgrades commenced after the initial LN work was completed, and therefore the DNAscope acquisitions were all acquired using the new upgraded system: the Zeiss Imager Z.2 microscope (TissueGnostics, Vienna, Austria).

### Multicolor flow cytometry (FACS)

Lymph node mononuclear cells (LMNCs) were characterized using multi-parameter flow cytometry analysis. Briefly, cells were stained with LIVE/DEAD Fixable Blue dead cell stain kit (Thermo Fisher Scientific, Waltham, MA, USA), CD3-BV711 (BD Biosciences, San Jose, CA), CD4-BV650 (BD Biosciences), CD19-PE-Cy5 (BioLegend, San Diego, CA, USA), HLA-DR-APC-CY7 (BioLegend) and, CD45-BV786 (BD Biosciences). For intracellular staining, cells were washed with PBS and incubated for 20 min with cytofix/cytoperm (BD Biosciences) according to manufacturer’s instructions. After fixation, cells were washed with perm wash buffer (BD Biosciences) and incubated for 20 min at RT with perm wash buffer containing CD68-AF488 (BioLegend) and CD206-PE (BD Biosciences) antibodies. Fluorescence minus one (FMO) or unstained cells were used as a control. Stained cells were acquired using the LSRFortessa (BD Biosciences) with FACSDiva™ software. Data were analyzed using FlowJo version 10.6.0 (FlowJo, LLC, Ashland, Oregon). LNMCs were identified as CD45^+^/CD3^-^/CD19^-^/HLA-DR^+^ by flow cytometry. Further gating was used to identify specific macrophage phenotypes as described in **(Fig S3)**.

### RNAscope *in situ* hybridization (ISH)

RNAscope ISH was conducted using the RNAscope 2.5 HD assay kit (Advanced Cell Diagnostics (ACD), Newark, CA, USA, Cat No: 322300) and the RNAscope multiplex fluorescent kit v2.0 (ACD, Cat No: 323100) as per manufacturer’s instructions. Lymph node FFPE blocks were sectioned into 5µm slices and affixed onto Surgipath X-tra adhesive pre-cleaned micro slides in preparation for antibody staining. Briefly, pre-treated samples were hybridized with the HIV-1 *gag-pol* probe (Cat No: 317691) at 40 °C for 2 hours. Next, the samples were incubated with signal amplification probes and horseradish peroxidase conjugated secondary antibodies. The signal was detected with Opal fluorophores (AKOYA Biosciences) for the multiplex fluorescent assay. Slides were imaged with Axio Observer and TissueFAXS imaging software (TissueGnostics).

### DNAscope *in situ* hybridization (ISH)

DNAscope ISH was conducted using the RNAscope 2.5 HD assay kit (Advanced Cell Diagnostics (ACD), Newark, CA, USA, Cat No: 322300) and the RNAscope multiplex fluorescent kit v2.0 (ACD, Cat No: 323100) as described for the RNAscope assay above. The following changes were made: the HIV-1-CladeC-*gag-pol*-sense (Cat number 444021) probe was used to detect HIV subtype C DNA and the slides were imaged using the Zeiss Imager Z.2 microscope (TissueGnostics, Vienna, Austria).

### Cell sorting and digital droplet PCR (ddPCR)

LNMCs were surface stained with the panel of antibodies including LIVE/DEAD fixable Aqua dead cell stain CD3 BV711, CD4 BV650, CD45 BV786, CD19 PE-CY5 and CD56 APC. Cells were sorted using the FACS Aria Fusion (BD Biosciences). Gating strategies for sorting are detailed in **(Fig S6)**.

Droplet digital PCR (ddPCR) was used to determine the HIV copy number in myeloid and paired CD4+ T cells from each study participant as previously described (84). DNA was extracted from myeloid and CD4+ T cells using DNeasy Blood & Tissue Kits (QIAGEN). Total HIV-1 DNA and host cell concentrations in the DNA extracts was estimated using primers and probes covering HIV-1 5′ LTR-gag, HXB2 coordinates 684– 810, (forward primer 5′-457 TCTCGACGCAGGACTCG-3′, reverse primer 5′-TACTGACGCTCTC GCACC-3′ probe/56-458 FAM/CTCTCTCCT/ZEN/TCTAGCCTC/ 31ABkFQ/), and human RPP30 gene (forward primer 5′-459 GATTTGGACCTGCGAGCG-3′, reverse primer 5′-GCGG CTGTCTCCACAAGT-3′, probe/56-460 FAM/CTGACCTGA/ZEN/AGGCTCT/31AbkFQ/). Thermocycling conditions were 95°C for 10 min, 45 cycles of 94 °C for 30 s and 60 °C for 1 min, 72 °C for 1 min. The Bio-Rad QX200 Droplet Reader was used to detect positively amplified DNA within droplets and data was analyzed using QX Manager Software v1.2 (Bio-Rad).

### Quantification and statistical analysis

#### High dimensional UMAP analysis

For dimensionality reduction and visualization, we employed the multiparametric global Uniform Manifold Approximation and Projection (UMAP) algorithm using R studio (version 2023.09.1+494), due to its effectiveness in revealing underlying structures in high-dimensional data. To identify distinct cell clusters within the UMAP space, we further employed the Flow Self-Organizing Map (FlowSOM) clustering algorithm, and to identify differential LNMC clusters. To further characterize the identified clusters, we utilized the unsupervised hierarchical clustering heatmap to summarize the mean fluorescence intensities of each loaded parameter, whereby warm colors (dark red) indicate high expression and cold colors (dark blue) indicate low expression. The frequency (%) was calculated and compared between cell clusters of interest to assess pro-inflammatory, anti-inflammatory and transitioning macrophage (85, 86). The frequency is calculated as follows:

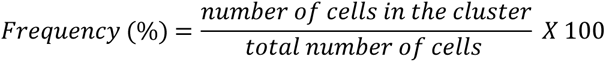

### Quantitative image analysis

#### Quantitative analysis of Lymph node macrophage frequency, and HIV Gagp24 protein co-expression by TissueQuest

Quantitative image analysis of macrophage frequency and HIV Gagp24 protein in whole tissue section scans was conducted with TissueQuest software (TissueGnostics). Total area measurements and nuclear segmentation analysis were performed on each whole tissue scan. The DAPI channel was used as a master channel for nuclei detection in each whole LN tissue section. Thereafter the grey scale images from each channel were analyzed and quantified according to their expression and reported as density/mm^2^ **(S2 Fig)**.

#### Quantitative analysis of HIV *gag-pol* DNA and RNA expression and CD68 co-expression by HALO

Quantitative image analysis of HIV vDNA and vRNA in LN tissues was performed using the FISH-IF module on HALO software (version v3.6.4134.396; Indica Labs) as previously described (87). Monochrome tiff image files of individual channels were exported from TissueFAXS and uploaded onto HALO where a merged image was formed and used for subsequent quantitative analysis. For the RNAscope analysis, GCs were segmented to form the ROIs for analysis. For the DNAscope analysis, GCs and EF regions which harbored high observed frequencies of vDNA were segmented to form the ROIs for analysis (on average, 1500 cells were segmented per ROI). 3 EF ROIs were segmented per donor and 1-3 GC ROIs were segmented per donor based on the frequency of GCs within the LN tissues. DAPI was used as a master channel to detect and segment all nuclei. Thereafter, thresholds were set to mask the vDNA or vRNA probe signals and CD68^+^ macrophages harboring vDNA/vRNA. The frequency of cells expressing HIV DNA or RNA was quantified as a proportion of DNA or RNA-harboring cells out of the total cells per region. The proportion of vDNA or vRNA arising from CD68^+^ macrophages was also assessed. The probe scoring system was set to routine parameters to reflect the minimum probe copies per cell summarized as probe score (minimum probe copies per cell): 0+(0); 1+(1); 2+(4); 3+ (10) and 4+(16).

The H-score indicates the amount of expression of the probe directed at gag-pol HIV DNA/RNA based on the minimum intensity thresholds. The H-score is calculated based on the equation:

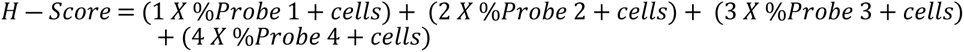

#### Quantification of Productively infected vs CD4-ingested CD68^+^ macrophages in lymph nodes

The percentage of productively HIV infected CD68^+^ macrophages vs. the percentage of CD4-ingested CD68^+^ macrophages was quantified manually. Intracellular CD4 staining within CD68^+^ macrophages was used to classify CD4-ingested macrophages. Membranous CD4 staining (in the absence of any intracellular CD4 staining) on CD68^+^ macrophages with intracellular Gagp24 positivity was used to classify productively infected macrophages. Briefly, two-four regions of interest were selected from HIV positive LN tissue sections (n=11) (**S6**). Thereafter, the average percentage (%) of productively infected CD68^+^ and CD4-ingested macrophages was computed.

### Statistical analysis

Statistical analysis and graphical presentation were performed using GraphPad Prism version 10.2.0 software (GraphPad Software Inc., La Jolla, CA, USA). For multi-group comparisons between 3 or more groups, the One-Way Analysis of Variance (ANOVA) was used for parametric datasets and the Kruskal Wallis test was used for non-parametric data comparisons. The Fisher’s exact test was used for demographic and clinical data comparisons involving ratios. The Shapiro-Wilk normality test was used to test for normality. The Student’s unpaired t test was used to compare differences between two unpaired groups with parametric data distribution. For non-parametric data distribution, the Mann-Whitney U test was utilized to compare differences between any two groups. For paired data analysis, the Student’s paired t test was used. For parametric data distribution, Pearson’s correlation was performed, and Spearman’s rank correlation analysis was performed for non-parametric data and the correlation rank (r) was reported as a measure of the strength of the association. A p value of <0.05 indicates statistical significance.

## Acknowledgements

We extend our appreciation to our study participants, the clinical and laboratory staff at FRESH and HPP. We acknowledge Dr. Thumbi Ndung’u (Africa Health Research Institute (AHRI), HPP, and Ragon Institute of MGH, MIT, and Harvard) and Dr. Bruce Walker (Ragon Institute of MGH, MIT, and Harvard) for their leadership in the FRESH cohort and for granting us access to samples from the FRESH and HPP Acute study cohorts. We acknowledge the Prince Mshiyeni Memorial Hospital and Durdoc Hospital. We would also like to thank the Optics and Imaging Core Lab at AHRI for using their facilities for tissue processing, embedding and sectioning. We acknowledge Dr. Jacob D Estes and the Estes Lab at the OHSU for DNAscope skills transfer and for generously providing us with control cell lines for assay validation. We also acknowledge Tatenda Chikowore (Ndung’u Lab-AHRI) for conducting the DNAscope probe sequence matching to our study cohort.

## Author contributions

Conceptualization, Z.M.N.; methodology, Z.M.N., M.M., C.C., T.K., B.M., and N.R.; investigation, M.M., C.C., T.K., B.M., S.M., N.R. and Z.M.N.; sample collection and processing, S.N., T.Ng., I.J., and J.P.; writing—original draft, M.M. and C.C.; writing—review & editing, Z.M.N., T.K., B.M., K.R. and N.R.; funding acquisition, Z.M.N.; resources, Z.M.N; supervision, Z.M.N., T.N., and K.R.

## Supporting information captions

**S1 Fig. Macrophage validation, phenotyping and localization. (A-D)** Representative multicolor IF images of: CD68 (white), CD3 (red) and KI67 (green) to define GCs **(A)**, CD68 (green), and IBA-1 (red) **(B)**, IBA-1 (red) from 2 LN tissue donors **(C)**, and the localization of CD68^+^ (white) macrophages relative to the LN GC indicated by BCL-6 (green) positivity from n=3 LN tissue donors **(D)**. The white dashed oval defines the GC. **(E)** Representative multicolor IF images from n=3 LN tissue donors of CD206^+^ (red) macrophages along LN lymphatic and blood vessels (white arrows). **(F)** Representative multicolor IF images from n=5 PLWH and n=5 PLWoH of CD68^+^ (white) and CD206^+^ (red) macrophages. Images acquired at 40x and all cell nuclei were stained with DAPI (blue). Scale bars= 100μm and 200μm. (TIF)

**S2 Fig. TissueQuest image analysis summary**. Summary of quantitative image analysis by TissueQuest. (TIF)

**S3 Fig. HIV modulates macrophage frequency. (A)** Multicolor IF images of the individual channels of CD206^+^ macrophages (red), CD68^+^ macrophages (white), and the composite image of CD68^+^CD206^+^ macrophages and nuclei counterstained with DAPI (blue) in LN tissue sections from n=2 representative PLWoH and n=2 representative PLWH. The white rectangles from the inner panels indicate regions of interest which are further magnified on the outer panels. The white circles indicate double positive CD68^+^CD206^+^ macrophages. Images were acquired at 40x. Scale bars= 100μm and 20μm. **(B)** Flow cytometry gating strategy of CD206^+^, CD68^+^ and CD68^+^CD206^+^ macrophages from LNMCs. (TIF)

**S4 Fig. HIV Gagp24 antibody controls and additional representative images from donors (A)** Representative Gagp24 negative control multicolor IF images from n=3 PLWoH of HIV Gagp24 antigen (brown) and KI67 (green) to define GCs. **(B)** Representative multicolor IF images from n=3 acute-treated and n=3 chronic-treated PLWH of GCs (white dashed ovals) indicated by BCL-6 (green) positivity and their respective expressions of CD68^+^ (white) macrophages and HIV Gagp24 antigen (brown). Nuclei are stained with DAPI (blue). Images acquired at 40x. Scale bars= 20μm, 50μm and 100μm. (TIF)

**S5 Fig. Representative RNAscope images from additional donors, H-scores and DNAscope validation. (A)** Representative images of FISH RNAscope coupled with multicolor IF of *gag-pol* HIV-1 RNA (red), CD68 (white) and KI67 (green) to define GCs in n=3 PLWH showing the individual channels and composite image. The white dashed oval defines the GC. Single RNA transcripts are shown as punctate dots. **(B)** Individual RNAscope probe H-scores per donor in n=4 PLWH quantified by HALO. Bars represent the mean and vertical lines represent standard deviation. **(C)** Alignment of DNAscope probe reference sequence to the sequences of our study cohort, validated with a 92.8% pairwise identity. **(D)** Representative DNAscope negative control images from n=2 PLWoH of *gag-pol* HIV DNA (red), CD68 (white) andKI67 (green). Cell nuclei were counterstained with DAPI (blue). Images acquired at 40x. Scale bars= 5μm and 100μm. (TIF)

**S6 Fig. Isolation of myeloid and CD4^+^ T cells from LNMCs and donor ratios of macrophage CD4-ingestion and productive infection. (A)** Flow cytometry gating strategy for CD4^+^ T cells and myeloid cells from LNMCs for ddPCR. **(B)** Individual donor ratios from n=11 PLWH of productively infected macrophages (green) and CD4-ingested macrophages (red) from high-resolution 100X oil immersion microscopy. (TIF)

**S1 Table. Participant demographics, clinical characteristics and experimental details.** (DOCX)

